# Tailored extracellular electron transfer pathways enhance the electroactivity of Escherichia coli

**DOI:** 10.1101/2021.08.28.458029

**Authors:** Mohammed Mouhib, Melania Reggente, Lin Li, Nils Schuergers, Ardemis A. Boghossian

## Abstract

Extracellular electron transfer (EET) engineering in *Escherichia coli* holds great potential for bioremediation, energy and electrosynthesis applications fueled by readily available organic substrates. Due to its vast metabolic capabilities and availability of synthetic biology tools to adapt strains to specific applications, *E. coli* is of advantage over native exoelectrogens, but limited in electron transfer rates. We enhanced EET in engineered strains through systematic expression of electron transfer pathways differing in cytochrome composition, localization and origin. While a hybrid pathway harboring components of an *E. coli* nitrate reductase and the Mtr complex from the exoelectrogen *Shewanella oneidensis* MR-1 enhanced EET, the highest efficiency was achieved by implementing the complete Mtr pathway from *S. oneidensis* MR1 in *E. coli*. We show periplasmic electron shuttling through overexpression of a small tetraheme cytochrome to be central to the electroactivity of this strain, leading to enhanced degradation of the pollutant methyl orange and significantly increased electrical current to graphite electrodes.

## Introduction

Bacteria are an inexhaustible natural resource and host to rich biochemical reaction networks, making them relevant for industrial applications across a wide range of sectors. To better tap into this metabolic potential, electron exchange with their environment is desirable. External energy input can be used as a driving force for anabolic reactions^[1,2]^. In contrast, catabolic reactions enable bioelectricity generation from organic substrates^[3]^, reductive extracellular synthesis of products such as nanoparticles^[4,5]^ and polymers^[6]^, as well as degradation of pollutants for bioremediation purposes^[7]^. In addition, quantification of the electron transfer can be exploited in biosensing applications^[8,9]^. Extracellular electron transfer (EET) is detrimental to the efficiency of these applications (Figure 1a).

**Fig. 1.**
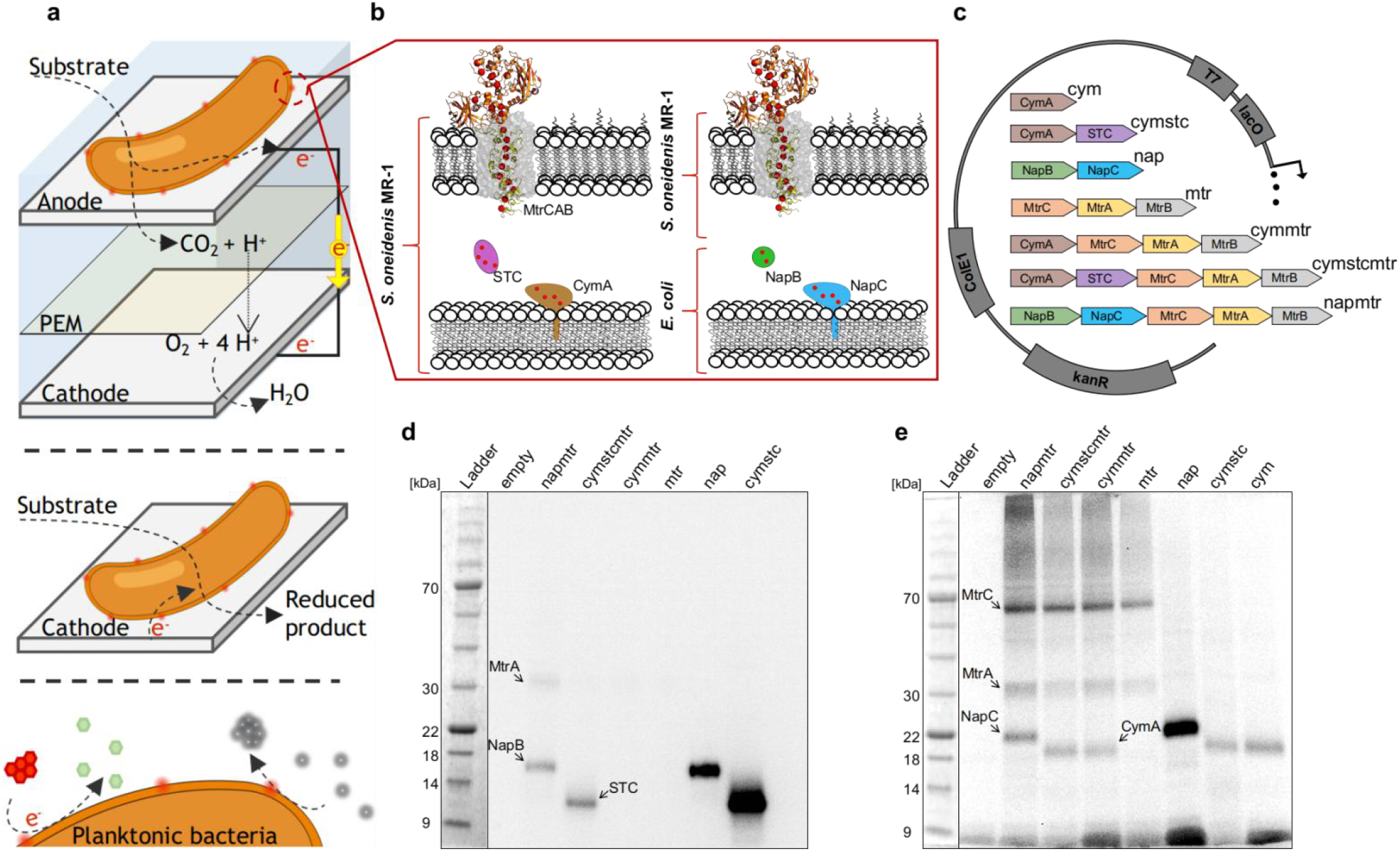
Engineering extracellular electron transfer pathways. (a) Schematic of a microbial fuel cell (top), a biocathode in microbial electrosynthesis (middle), and electrode free applications of EET in bio-synthesis and -remediation (bottom). (b) Overview of the S. *oneidensis* MR-1 and *E. coli* cytochromes used in this study and their localization. (c) Constructs for the expression of engineered EET pathway variants and strain names. (d) Size separated cytochromes in the periplasm and (e) membrane fraction of engineered strains. SDS-PAGE (d, e) was performed using 30 μg of protein per lane, 4-20 % polyacrylamide gels and MOPS running buffer. Cytochromes were detected using enhanced chemiluminescence.

Exoelectrogens are bacteria that evolved EET pathways enabling the use of extracellular compounds as terminal electron acceptors^[3,10]^. EET occurs through different direct and mediated mechanisms^[11–15]^, with one of the best characterized pathways being the metal-reducing (Mtr) pathway from *Shewanella oneidensis* MR-1^[15]^. Here, electrons are transported across both insulating cellular membranes and the periplasmic space through closely stacked heme cofactors in c-type cytochromes. The inner membrane cytochrome CymA can oxidize quinol for electron transfer to various periplasmic cytochromes, which pass the electron to the outer membrane-spanning Mtr complex and thus to the extracellular space^[15]^. While exoelectrogens are capable of efficient EET, they often lack in characteristics desirable for application in microbial electrochemical systems (MES), such as a broad substrate spectrum and ease of metabolic engineering for specific product syntheses. On the contrary, *Escherichia coli* lacks in EET capabilities but would be suitable for a wide range of applications given its metabolic flexibility and readily available synthetic biology tools from decades of research.

*E. coli* was previously engineered for EET through heterologous expression of parts of the Mtr pathway^[16–21]^. The expressed electron transfer pathways differ from study to study, ranging from the expression of single cytochromes^[19,20]^ to simultaneous expression of cytochromes in both cellular membranes^[18]^ to co-expression of CymA in the inner membrane with periplasmic cytochromes^[17]^. However, EET efficiencies remain low, with open questions remaining as to which cytochromes are essential for EET in engineered *E. coli*.

For instance, the Mtr complex can be omitted in the presence of an extracellular redox mediator capable of diffusing into the periplasm^[17]^. This reduces the metabolic cost of expressing the EET pathway, as the Mtr complex necessitates post translational modification for the incorporation of 20 heme cofactors^[11, 22]^. However, direct electron transfer would be impeded, and mediated transfer could potentially be facilitated through interaction of the redox mediator with MtrC.

Direct transfer of electrons from CymA to MtrA is possible^[23]^, and *E. coli* expressing only membrane-associated cytochromes while omitting periplasmic shuttles were shown to be capable of EET^[18]^. However, recent structural insights into the Mtr complex suggest that MtrA is anchored in the Mtr complex with its heme closest to the N-terminus protruding only 2 nm into the periplasmic space^[11]^. In addition, studies on electron transfer in *S. oneidensis* MR-1 highlight the importance of periplasmic cytochromes in EET ^[24, 25]^. Given the similar width of the periplasmic space in *E. coli (~20 - 30 nm)*^[26,27]^ and *S. oneidensis* MR-1 (~23.5 nm)^[28]^, these findings suggest that co-expression of periplasmic cytochromes may enhance EET in engineered *E. coli*.

While *E. coli* are not natively capable of respiration on extracellular solids, organic compounds can act as terminal electron acceptors in the periplasm under anaerobic conditions^[29]^. Overexpression of the herin involved cytochromes may be beneficial to EET in *E. coli*, especially in conjuncture with heterologously expressed cytochromes to enable further electron transfer across the outer membrane.

Given emerging applications of MES and their fundamental reliance on efficient EET, we aimed to construct an optimized EET pathway in *E. coli*. To this end, we expressed different EET pathways using components from both *S. oneidensis* MR-1 and *E. coli* distributed across the periplasmic space, and both membranes. Contributions to electron transfer were elucidated using colorimetric electron transfer assays based on the reduction of methyl orange and ferric citrate as soluble electron acceptors.

Although native *E. coli* cytochromes could enhance EET, electron transfer rates lagged behind their *Shewanella oneidensis* MR-1 counterparts. We found the small tetraheme cytochrome (STC) to significantly improve electron transfer rates, with cooperative effects when coexpressing the Mtr complex, highlighting the importance of periplasmic shuttles in EET pathway design. Using the EET pathway with the highest electron transfer rate to soluble electron acceptors in a lactate fueled electrochemical cell, we observed increased current generation as compared to a control strain lacking heterolougous expression of cytochromes, and the same strain lacking a periplasmic electron shuttle. We envisage this final engineered strain to serve as a tool for us and others to enhance the efficiency of existing MES, as well as for the development of new MES with an extended range of products and substrates.

## Results and discussion

To enhance EET in *E. coli*, we used two design approaches for heterologous expression of electron transfer pathways (Figure 1b). Firstly, we sought to express the Mtr pathway from *S. oneidensis* Mr-1, including a soluble cytochrome for electron shuttling across the periplasmic space. To this end, we expressed CymA in the inner membrane, the Mtr complex in the outer membrane, and the soluble cytochrome STC in the periplasm. While *S.oneidensis* MR-1 harbors a large pool of cytochromes involved in electron transfer across the periplasm, STC was selected due to its role in electron transfer between CymA and MtrA^[24,30]^. In addition, heterologous expression of STC alongside CymA and MtrA was previously shown to improve reduction of a soluble electron acceptor, as compared to expression of other periplasmic cytochromes from *S. oneidensis* MR-1^[17]^. As electron transfer through CymA may be limiting electron transfer rates in engineered *E. coli*^[31]^, its oxidation by STC for rapid turnover makes this approach especially attractive to increase EET. In a second approach, we aimed to combine cytochromes from both *E. coli* and *S. oneidensis* MR-1 to leverage the native ability of *E. coli* for electron delivery to the periplasm in a hybrid EET pathway. Here, expression of CymA in the inner membrane was circumvented by expression of the *E. coli* cytochromes NapC and NapB involved in periplasmic reduction of nitrate. *E. coli* is also able to reduce nitrite and trimethylamine-N-oxide in the periplasm using the nrf and TMAO reductase systems, respectively ^[29]^. However, the nitrate reductase gene cluster is of particular interest in constructing a hybrid EET pathway. Its inner membrane component NapC and CymA are similar in both sequence and function^[20]^. In addition, a homologue to its periplasmic component NapB may play a role in EET through the Mtr pathway in *S. oneidensis* MR-1^[32]^. While there is a genomic copy of both cytochromes present in *E. coli*, they are only expressed under anaerobic conditions and further induced in the presence of nitrate, as nitrate is not the preffered terminal electron acceptor in *E. coli*^[29, 33]^.

In addition to the two complete pathways incorporating inner membrane, outer membrane and periplasmic cytochromes, five partial pathways were expressed to evaluate the importance of different cytochromes in electron transfer and their potential omission for a minimal EET pathway. All genes constituting the electron transfer pathways were expressed in single operons under the control of an IPTG inducible promoter (Figure 1c, Table S1). Alongside cytochromes for the EET pathway, the cytochrome c maturation (ccm) machinery necessary for posttranslational incorporation of hemes was expressed constitutively using a tet promoter on a separate plasmid. Although there is a genomic copy of the ccm machinery, it is not expressed under aerobic conditions.

Protein samples from all strains (names specified in Figure 1c) expressing electron transfer pathways and a control strain harboring an empty vector backbone (“empty”) were analyzed to verify cytochrome expression levels and localization. Periplasmic and membrane fractions were separated using SDS-PAGE. Gels were directly stained for cytochromes (Figures 1d and e) and subsequently for their total protein content (Figure S1). The latter revealed that heterologously expressed cytochromes constitute a comparatively low fraction of periplasmic and membrane proteins. STC forms an exception, as its high expression levels allowed for detection together with other periplasmic proteins using coomassie brilliant blue.

Both STC and NapB were localized in the periplasm, with much higher expression levels in strains where the Mtr complex was omitted from the pathway. We hypothesize that this may either be caused by decreased mRNA stability and secondary structures in longer transcripts including the *mtrCAB* genes, or competition of multiple co-expressed cytochromes for post translational modification. Interestingly, MtrA was detected in the periplasm and membrane fraction, while MtrC was only observed in the latter. This shows that the MtrA band in the periplasm does not stem from membrane fragments in the periplasmic fraction, and is in agreement with previous reports on its localization^[16,19]^. The cytochromes MtrC, MtrA, CymA, and NapC were all localized in the membrane fraction as expected.

To study the efficiencies of the different electron transfer pathways, we first compared electron transfer rates to soluble electron acceptors using ferric citrate and methyl orange (Figure 2a). Methyl orange is a suitable substrate for colorimetric EET assays, as reduction of the azo group is accompanied by a loss of absorbance at 420 nm^[34]^. While its reduction occurs predominantly extracellularly, interaction with intracellular components is possible through redox mediators and potentialy limited uptake^[35]^. Methyl orange was used as the sole electron acceptor under anaerobic conditions, with glycerol as the electron donor in a minimal growth medium (M9). Both the concentration of methyl orange and the optical density of cultures were monitored over time (Figure S2). Over two days, we observed a decrease of the methyl orange concentration, which could be approximated by a pseudo first-order decay reaction (Figure S2). Experiments were carried out using 12 biological replicates per strain, and the resulting rate constants were normalized to an OD_600_ of 1 (Figure 2b). Differences in the mean methyl orange reduction rates were compared using two-tailed T-tests, and the resulting probabilities are summarized in Figure S3.

**Fig. 2.**
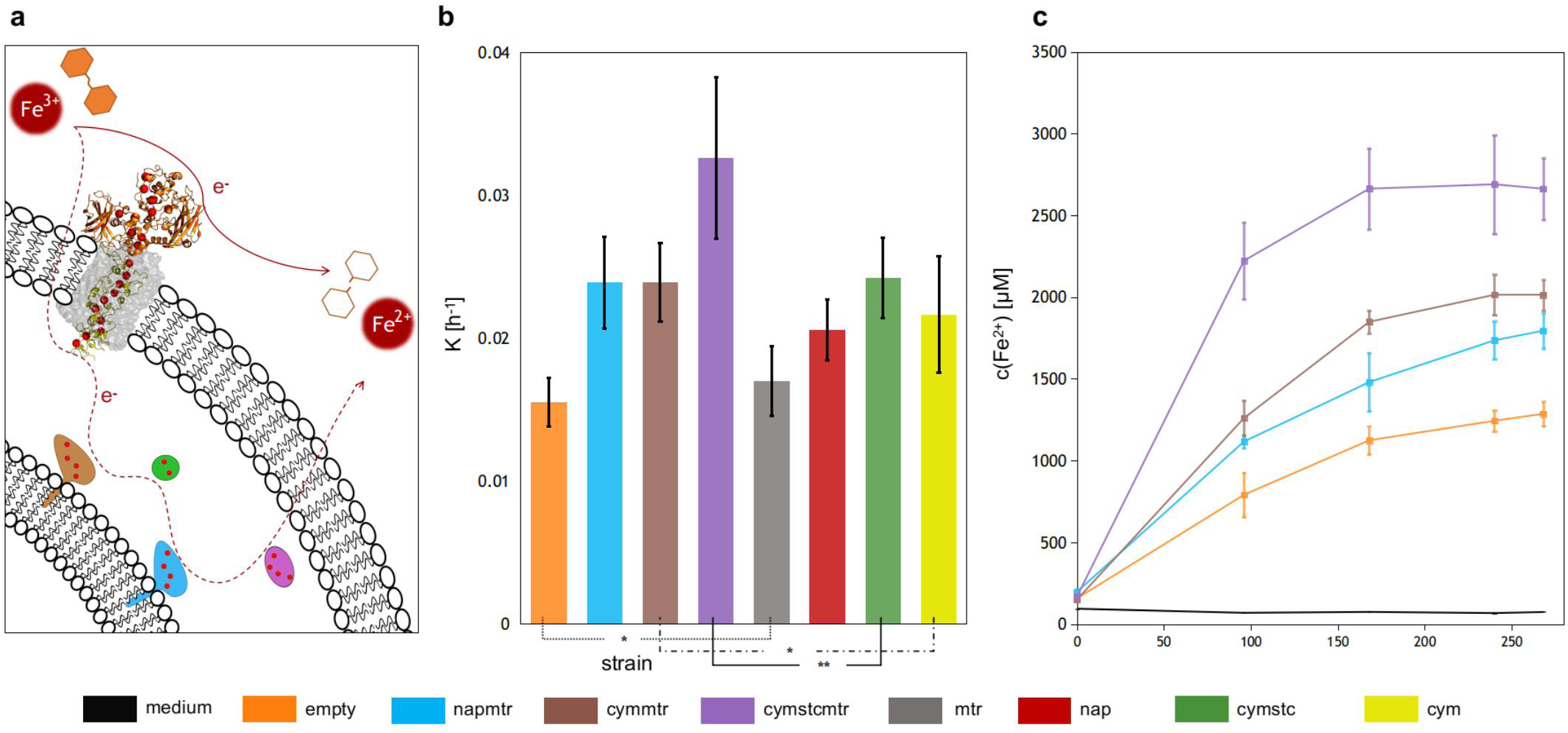
Election transfer assays using soluble electron acceptors. (a) Schematic illustration of methyl orange and ferric citrate reduction by engineered *E. coli*. (b) Reaction rate coefficients for reduction of methyl orange by engineered *E.coli* at an OD_600_ of 1. (c) Ferri citrate reduction over 11 days monitored using Fe^2+^ accumulation in assay media. Both reduction assays were carried out in minimal media with the respective electron acceptor under anaerobic conditions. *(p>0.15) **(p<0.01)

With exception of the mtr *E. coli* (p=0.11), all EET pathways significantly increased methyl orange reduction rates (Figures 2b, S3) compared to the empty vector control strain, suggesting reduction of methyl orange occurs via direct or indirect interaction with expressed cytochromes. This includes the *E. coli* cytochromes NapB and NapC, which led to reduction rates on par with the cymmtr *E. coli* when coexpressed with the Mtr complex (napmtr), showing that it is possible to substitute CymA by *E. coli* native cytochromes. However, the complete Mtr pathway, including STC (cymstcmtr), showed by far the highest reduction rate, which differed significantly from all other strains (Figure S3).

Interestingly, while expression of the Mtr complex on its own and alongside CymA leads to an insignificant increase in reduction rates, its contribution to electron transfer rates increases when other cytochromes are coexpressed in the inner membrane and periplasm (Table 1). This shows that electron delivery across the inner membrane and the periplasm is limiting in this strain, and can not be solely substituted by potential *E. coli* native mechanisms. Analogous to the Mtr complex, expression of STC alongside CymA (cymstc) does not increase reduction rates significantly beyond the expression of CymA alone. However, STC contributes significantly to reduction rates when coexpressed with CymA and the Mtr complex (cymstcmtr). This positive cooperative effect on electron transfer by co-expression of STC and the Mtr complex suggests interaction of these cytochromes in engineered *E. coli*. Moreover, it indicates periplasmic shuttling to be crucial for the involvement of the Mtr complex in the reduction reaction and thus for higher electron transfer rates. The importance of this interaction is further underlined by much higher STC expression in the cymstc strain as compared to the cymstcmtr strain (Figures 1d, S1b), coupled with significantly lower reduction rates. Enhanced electron delivery to MtrC may be increasing methyl orange reduction rates by promoting reduction at the outer membrane, and thus avoiding diffusion limits.

**Tab. 1.** Electron transfer assays using soluble electron acceptors. Relative contribution of Mtr complex or STC coexpression in different strains to reduction rates.

| reference strain | $\frac{K_{ref + mtr/stc} - K_{ref}}{K_{ref}}$ | CI<sub>95%</sub> | *p* |
| --- | --- | --- | --- |
| MtrCAB + |  |  |  |
| empty | 9.4 % | ± 10.2 % | 0.11 |
| cym | 10.5 % | ± 12.8 % | 0.12 |
| cymstc | 34.7 % | ± 16.0 % | 0.0005 |
| STC + |  |  |  |
| cym | 11.9 % | ± 13.8 % | 0.10 |
| cymmtr | 36.4 % | ± 14.9 % | 0.0004 |

To assess whether we observe methyl orange specific effects or electron transfer efficiencies in general, we also performed electron transfer assays with ferric citrate. Ferric citrate could diffuse into the periplasm^[36]^, and the reduction could be monitored using a colorimetric assay based on the complexation of Fe^2+^ with ferrozine^[37]^. We monitored the Fe^2+^ concentration and the OD_600_ over eleven days under anaerobic conditions, with glycerol as the sole carbon source (Figures 2c, S4). Both of the complete EET pathways (cymstcmtr, napmtr), the Mtr pathway lacking periplasmic shuttling (cymmtr), and the empty vector control were studied using three biological replicates. Reduction of Fe^3+^ followed the same trend as methyl orange reduction, with the cymstcmtr strain showing the highest reduction over time, all cytochromes showing higher reduction than the empty vector control, and the hybrid EET pathway incorporating NapB and NapC exhibit a similar electron transfer rate as compared to the cymmtr strain. The increased reduction rates for two distinct soluble electron acceptors by the cymstcmtr strain clearly show that periplasmic shuttling promotes electron transfer in engineered *E. coli*.

Efficient EET to electrodes is crucial for microbial electrochemical systems (Figure 1a). Therefore we used the cymstcmtr strain in chronoamperometric (CA) measurements with a graphite felt electrode as the electron acceptor and lactate as the electron donor (Figure 3a). Bioelectricity generation was compared to the empty vector control and the cymmtr strain to assess whether co-expression of STC improves EET. The experiments were carried out at a positive applied potential of 0.2 V against an Ag/AgCl reference electrode, with a platinum wire as the counter electrode. After three days the system reached a steady state, with the cymstcmtr *E. coli* showing an approximately 2- and 3-fold higher current compared to the cymmtr and control strain, respectively.

**Fig. 3.**
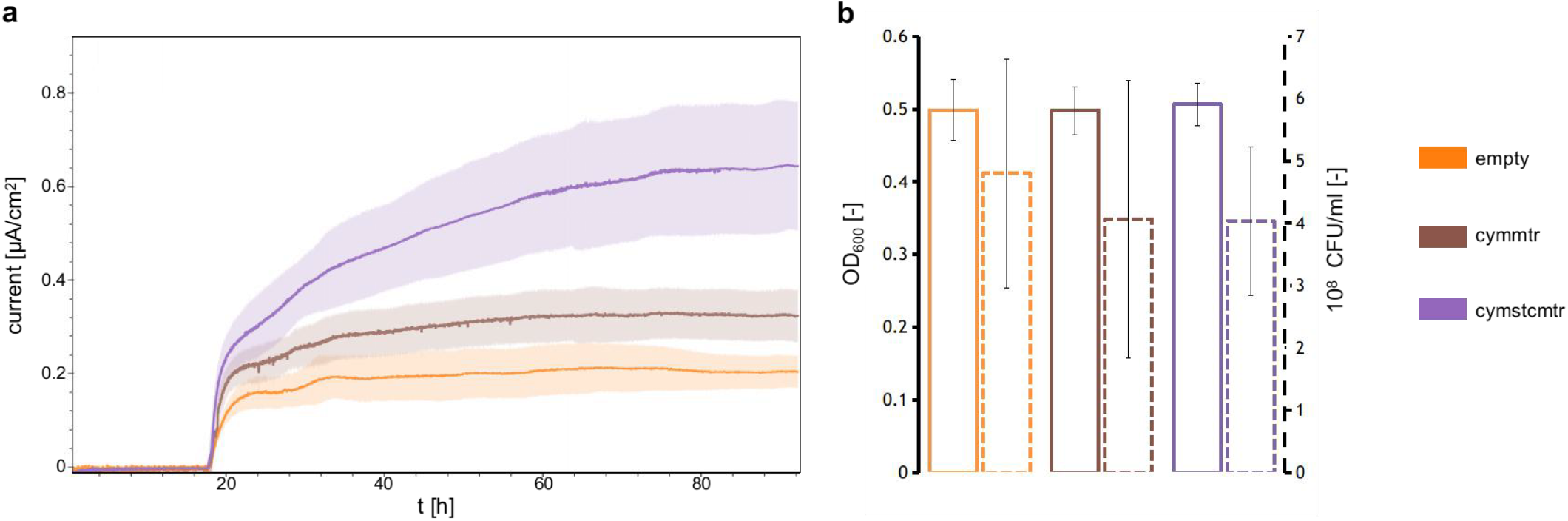
Electron transfer to graphite as the extracellular electron acceptor. (a) Chronoamperometric measurments in a lactate fueled electrochemical cell. (b) Measurments of the OD_600_ in the electrolyte and assessment of relative electrode colonization using CFU counts from electrode extracts after chronoamperometric measurments.

This increase in current suggests that the cymstcmstr strain may be applied in bioelectrochemical systems for improved device efficiencies. Since the higher current may not stem from higher EET rates, but could reflect enhanced survival of the cymstcmtr *E. coli* under electrode reducing conditions, measurements of the OD_600_ from the electrolyte after CA measurements were taken (Figure 3b). This revealed no significant differences in growth between strains. In addition, there was no preferred colonization of the electrode by any of the strains, as assessed through estimation of colony-forming units (CFU) on working electrodes (Figure 3b). As the increase in current from the cymstcmtr strain could stem from changing metabolic activity, we evaluated the lactate consumption and formation of different fermentation products in electrochemical cells (Figure S5). These measurements revealed no significant differences between strains, with lactate consumption after three days showing that increase in current from cytochrome expressing strains is not caused by increased catabolic activity.

## Conclusion

After evaluation of different EET pathways in *E. coli*, expression of the Mtr pathway including the periplasmic shuttle STC yielded in the highest electron transfer rates to both soluble and solid electron acceptors. This strain (cymstcmtr) constitutes a complete implementation of the Mtr pathway in *E. coli*, for a total of four heterologously expressed cytochromes with 28 heme cofactors distributed across the periplasm and both cellular membranes. Positive cooperative effects between expression of the Mtr complex and STC on the reduction of methyl orange, and the higher reduction rates for methyl orange, ferric citrate and graphite felt electrodes show periplasmic shuttling to be essential for EET in engineered *E. coli.* This finding is in agreement with previous studies of the Mtr pathway in *S. oneidensis* MR-1^[24, 25]^. Increased electron transfer rates compared to expression of NapBC from *E. coli* alongside the Mtr complex (napmtr), may be due to the Mtr pathway undergoing natural evolution in metal respiring bacteria. Protein engineering approaches such as directed evolution could potentially improve non native interactions between NapB and MtrA, or alternatively NapC and STC.

The postive effect of STC expression on EET confirms that electron delivery across CymA in the inner membrane was limiting EET^[31]^. Since reduction of STC improves turnover of CymA, the system was limited in electron transfer from CymA to MtrA. Future bioengineering approaches to enhance EET could therefore target accumulation of NADH and ubiquinol instead of the Mtr pathway, to increase electron delivery to the inner membrane. Modification of the ccm machinery was previously shown to increase CymA expression levels^[31]^, and could be coupled with the here reported improved electron transfer pathway to further enhance the electroactivity of engineered *E. coli*.

The enhanced degradation of methyl orange, and the increased current to graphite electrodes demonstrates the utility of the cymstcmtr strain for application in bioremediation and energy applications. As *E. coli* is relatively easy to engineer, this strain could be adapted to specific applications, such as targeted degradation of defined feed substrate compositions for improved performance of microbial fuel cells. The herein presented engineered *E. coli* has the potential to improve device efficiencies in existing *E. coli* based MES, to serve in the development of new microbial electrochemical systems, and to be used as a reference in future optimizations of EET in E. coli.

## Experimental section

### Plasmid preparation

All strains, plasmids and primers can be found in table S1. *E. coli* DH5α was used for all molecular cloning experiments. Plasmids pMO1 (napBC) and pMO2 (cymA, cctA) were procured from GenScript using the pSB1ET2 plasmid as the backbone, with the coding sequence for NapBC from *E. coli* K12 MG1655 (genebank: CP032679.1), and for CymA and STC from *S. oneidensis* MR-1 (genebank: AE014299.2). The MtrCAB coding gene was amplified from plasmid I5049 using primers 1, 2 or 3, 4 for gibson assembly^[38]^ with pMO1 (primers 5, 6) and pMO2 (primers 7, 8) respectively, yielding plasmids pMO1.1 and pMO2.1. For construction of pMO2.2, I5049 was cut (EcoRI and XbaI) for excision of the MtrCAB coding sequence, blunted and religated.

### Bacterial growth

E. *coli* C43(DE3) were transformed with pEC86 and either pSB1ET2, pMO1, pMO1.1, I5023, pMO2, pMO2.1, pMO2.2 or I5049 using electroporation, and plated on 2xYT-agar plates with kanamycin and chloramphenicol. Colonies were used to inoculate 5 ml 2xYT (kan, cam) medium, and grown overnight (220 rpm, 37°C). Expression cultures (2xYT, kan, cam) were inoculated to an OD_600_ of 0.1, and grown until an OD_600_ of ~0.6 (220 rpm, 37°C). Cytochrome expression was then induced using 10 μM IPTG, and cultures were grown overnight (220 rpm, 30°C, 16 h).

### Subcellular fractionation

Subcellular fractionation was adapted from Malherbe et. al.^[39]^. In brief, cytochrome expression cultures were harvested (3000 g, 4°C, 15 min) to an OD_600_ of 8 in 16 ml. Cells were washed in PBS (3000 g, 4°C, 15 min), resuspended in buffer A (100 mM Tris-HCl pH 8, 500 mM succrose, 0.5 mM EDTA), incubated for 5 minutes on ice and centrifuged (3000 g, 4°C, 15 min). For recovery of periplasmic proteins pellets were resuspended in 1 mM MgCl_2_ and incubated for 2 minutes. After centrifugation (3000 g, 4°C, 15 min), supernatants containing periplasmic proteins were concentrated using 3 kDa molecular weight cutoff Amicon filters.

Pellets were washed once in buffer B (50 mM Tris-HCl pH 8, 250 mM succrose, 10 mM MgSO_4_), resuspended in buffer C (50 mM Tris-HCl pH 8, 2.5 mM EDTA) and lysed through sonication. Lysates were cleared by centrifugation (10000 g, 10 min, 4°C), and membrane proteins were recovered by several centrifugation (21000 g, 4 h, 4°C) and washing steps, with final resuspension of pellets in buffer C with 5 % Triton-X100 at 4°C.

### In-gel cytochrome staining

Proteins were seperated according to size using SDS-PAGE (SurePage 4%-20% gels, 1x NuPAGE LDS sample buffer, MOPS running buffer).

Polyacrylamide gels were rinsed in distilled water, and excess water was removed before application of an ECL stain (SuperSignal West Pico PLUS, Thermo Scientific) to the gel followed by incubation for 5 minutes. Gels were imaged for cytochromes using an ECL imaging system (Fusion solo S, Vilber).

### Methyl orange

Cytochrome expressing *E. coli* were harvested (3000 g, 4°C, 15 min), washed in M9-glycerol medium^[17]^, and resuspended in 10 ml M9 glycerol medium with 200 μM methyl orange to an OD_600_ of 4. Cultures were sealed for anaerobic growth and stirred at room temperature for 3 days. Culture samples were taken with a syringue at set time intervals and the OD_600_ was montiored using a spectrophotometer (UV-3600 Plus, Shimadzu). After centrifugation of samples (22°C, 21000g, 3 min), clear supernatants were used for absorbance measurements at 465 nm in 96 well plates (Varioscan LUX, Thermo Scientific).

### Ferric citrate reduction assay

Cytochrome expressing *E. coli* were harvested (3000 g, 4°C, 15 min), washed in M9 lactate medium^[16]^, and resuspended in 10 ml M9 lactate medium with 10 mM Fe(III)-citrate to an OD_600_ of 2. Cultures were sealed for anaerobic growth and stirred at room temperature for 10 days. Culture samples were taken with a syringue at set time intervals and the OD_600_ was montiored using a spectrophotometer (UV-3600 Plus, Shimadzu). Samples were filtered (0.2 μM) and diluted to 50 % in 1 M HCl. Absorbance was measured at 562 nm in 96 well plates, immediatly after addition of 180 μl 1 M ferrozine solution to 20 μl of dilute samples.

### Electrochemical setup and measurments

Electrochemical measurments were carried out in a single chambered cell with M9-lactate medium^[40]^ as the electrolyte, a Pt wire counter electrode, an Ag/AgCl reference electrode and a 1 × 1.1 × 0.5 cm graphite felt (Alfa Aesar) electrode connected to a Pt wire as the anode. Anodes were pretreated in piranha solution (97% H2SO4: 30% H2O2 in 3:1 mixture) for 10 minutes, before washing and storage in distilled water. Experiments were run using a multichannel potentiostat (MultiPalmSens 4).

Bacteria were harvested (3000 g, 4°C, 20 min) and washed 3 times in M9 medium with carbon sources as indicated. Final pellets were resuspended in 5 ml of the same medium and used to inoculate electrolytes for a final volume of 100 ml. Electrolytes were supplemented with kanamycin, chloramphenicol and 10 μM IPTG.

After measurements, anodes were rinsed three times and incubated in PBS (2h, 4°C, 15 ml) for characterization of bacterial growth using CFU counts.

### HPLC analysis

Samples from electrochemical cells were filtered (0.2 μM) and seperated on a Hiplex H column (Agilent) using 14 mM H_2_SO_4_ as the mobile phase. The column temperature was kept constant at 40°C. Experiments were carried out on an Infinity II (Agilent) HPLC system, and compounds were detected using absorbance (210 nm) and refractive index measurments (Agilent 1260 DAD WR, 1260 RID)

## Acknowledgements

The authors are grateful for funding by the Gebert-Rüf Stiftung.

