## Supplementary_Information for "Tailored extracellular electron transfer pathways enhance the electroactivity of Escherichia coli"

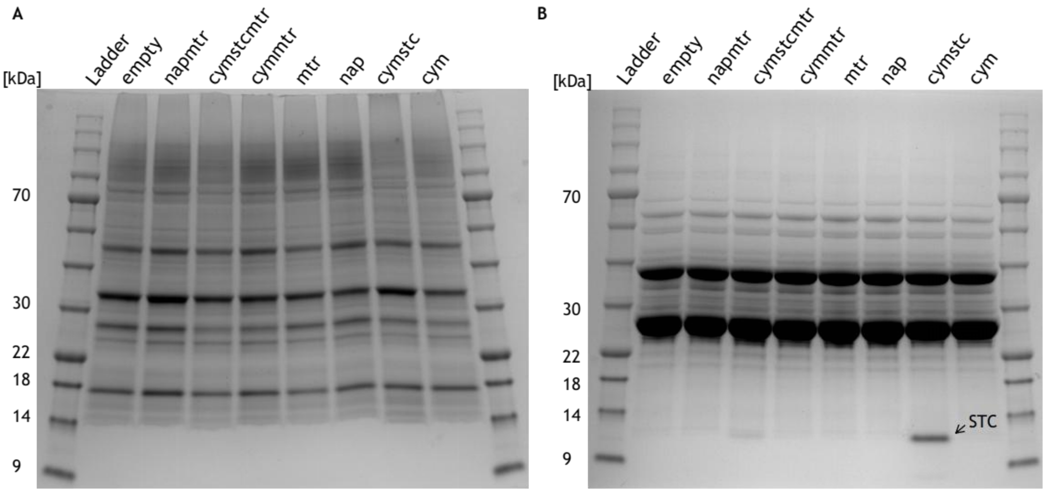


**Fig. S1 Coomassie Brilliant blue G250 staining of SDS-PAGE gels** from (A) periplasmic and (B) membrane fractions after prior enhanced chemiluminescence staining for cytochromes (see Figure 1d-e in the main text)

**Tab. S1 Overview of bacterial strains, plasmids and primers used in this study.**

| Strains, plasmids and primers | Characteristics | Source |
|---|---|---|
| **Strains** | | |
| *E. coli* DH5α | | |
| *E. coli* C43 (DE3) | | |
| *S. oneidensis* MR-1 | | |
| **Plasmids** | | |
| pEC86 | CcmABCDEFGH expression, chloramphenicol resistance, tet promoter | [41] |
| pSB1ET2 | empty backbone for cytochrome expression, kanamycin resistance, T7 promoter | [16] |
| pMO1 | NapBC expression, kanamycin resistance, T7 promoter | This study |
| pMO1.1 | NapBC and MtrCAB expression, kanamycin resistance, T7 promoter | This study |
| I5023 | MtrCAB expression, kanamycin resistance, T7 promoter | [16] |
| pMO2 | CymA and STC (cctA gene) expression, kanamycin resistance, T7 promoter | This study |
| pMO2.1 | CymA, STC and MtrCAB expression, kanamycin resistance, T7 promoter | This study |
| I5049 | CymA and MtrCAB expression, kanamycin resistance, T7 promoter | [18] |
| pMO2.2 | CymA expression, kanamycin resistance, T7 promoter | This study |
| **Primers** | | |
| 1 | 5`- -3` gtttttaacagggttattgcgaattctatttccagcatccacttaagt | |
| 2 | 5`- -3` ctgcagcggccgctactagttctagattagagtttgtaactcatgctca | |
| 3 | 5`- -3` agaagtaattgccgctgcaaGaattctatttccagcatccacttaagtc | |
| 4 | 5`- -3` ctgcagcggccgctactagttctag | |
| 5 | 5`- -3` gttacaaactctaatctagaactagtagcggccgctgcag | |
| 6 | 5`- -3` ggatgctggaaatagaattcgcaataaccctgttaaaaacctggc | |
| 7 | 5`- -3` gttacaaactctaatctagaactagtagcggccgctgcag | |
| 8 | 5`- -3` ggatgctggaaatagaattCttgcagcggcaattacttcttc | |

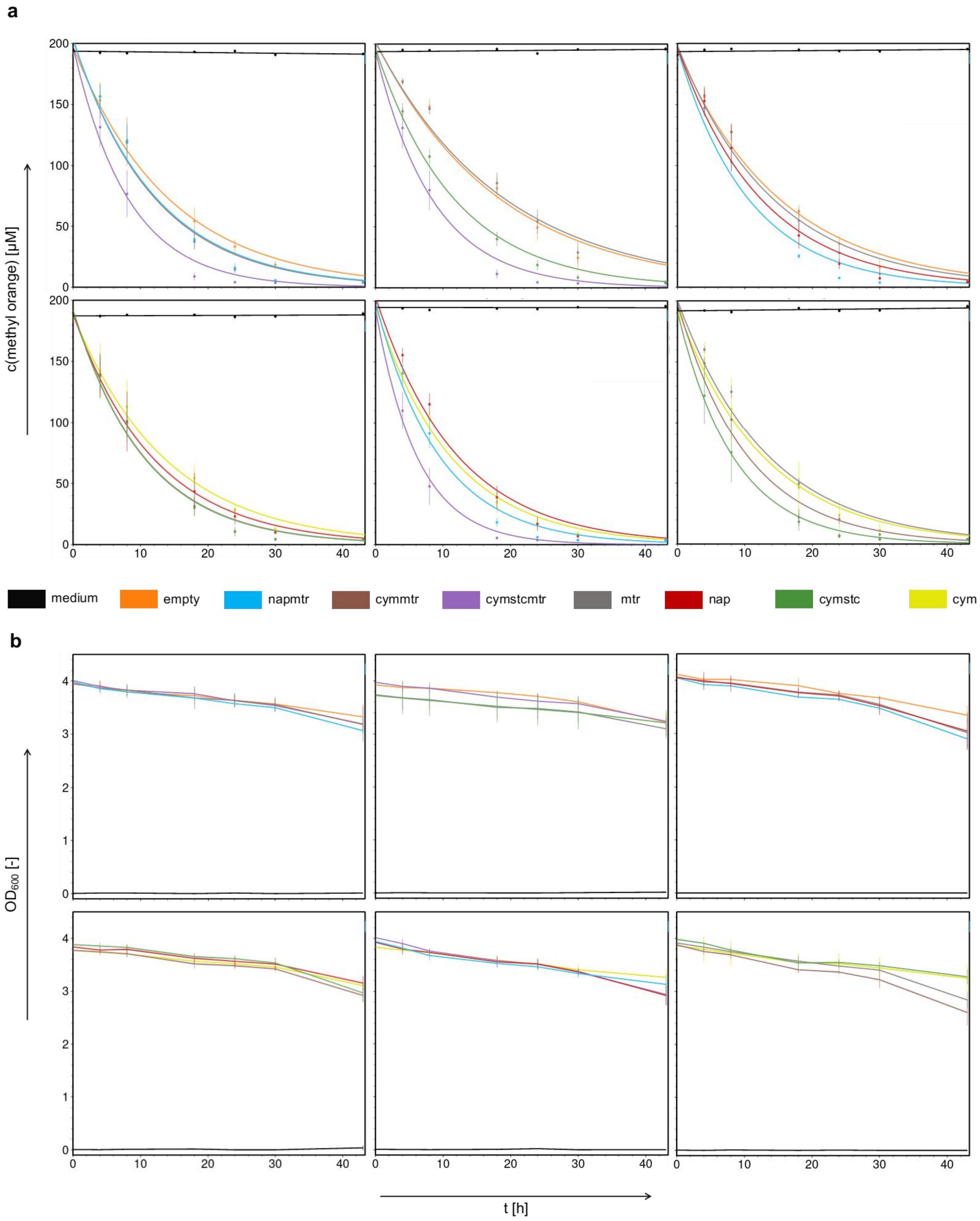


**Fig. S2 Methyl orange reduction assay.** Measurments of the (a) methyl orange concentration and (b) $OD_{600}$ in assay tubes over the course of 2 days.

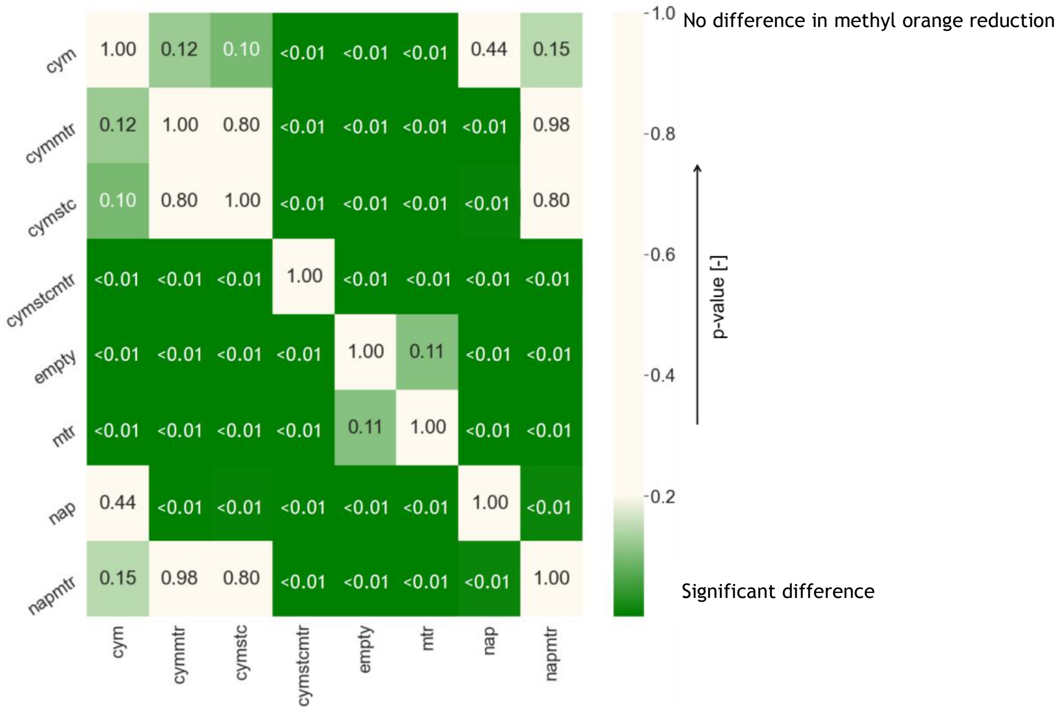


**Fig. S3 Significance evaluation of observed differences in mean methyl orange reduction rates.** P values obtained from two tailed two sample T-tests without assuming equal population variance (Welch´s T-test) plotted against each strain pair.

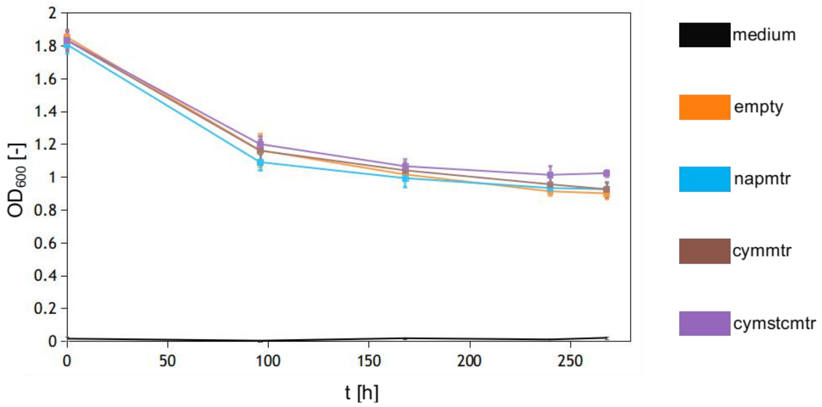


**Fig. S4.** Measurments of the $OD_{600}$ over the course of 11 days during ferric citrate reduction assays.

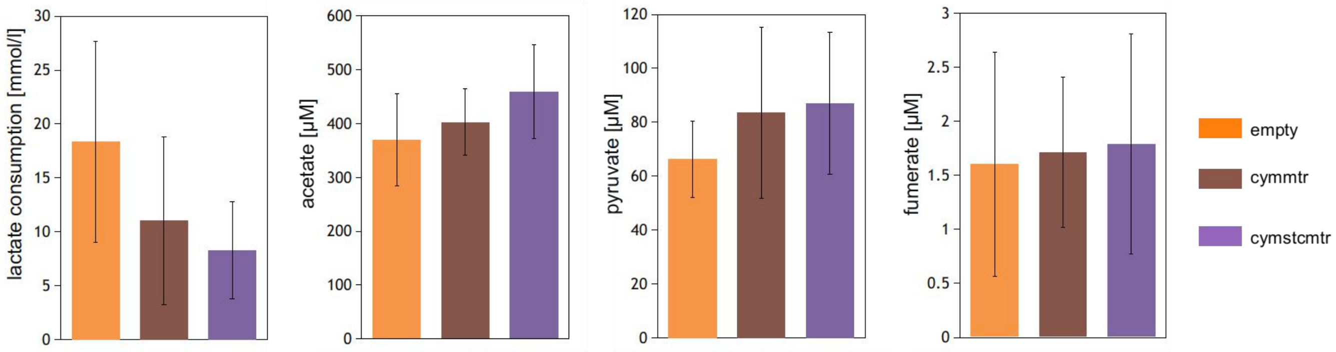


**Fig. S5.** Lactate consumption and concentration of different fermentation products after chronoamperometric measurments in lactate fueled electrochemical cells.
